# A novel mechanism for the loss of mRNA activity in lipid nanoparticle delivery systems

**DOI:** 10.1101/2021.09.21.461221

**Authors:** Meredith Packer, Dipendra Gyawali, Ravikiran Yerabolu, Joseph Schariter, Phil White

## Abstract

Lipid nanoparticle (LNP)-formulated mRNA vaccines were rapidly developed and deployed in response to the SARS-CoV-2 pandemic. Due to the labile nature of mRNA, identifying impurities that could affect product stability and efficacy is crucial to the long-term use of nucleic-acid based medicines. Herein reversed phase ion pair high performance liquid chromatography (RP-IP HPLC) was used to identify a class of impurity formed through lipid:mRNA reactions; such reactions are typically undetectable by traditional mRNA purity analytical techniques. The identified modifications render the mRNA untranslatable, leading to loss of protein expression. Specifically, an electrophilic impurity derived from the ionizable cationic lipid component is shown to be responsible. Mechanisms implicated in the formation of reactive species include oxidation and subsequent hydrolysis of the tertiary amine. It thus remains critical to ensure robust analytical methods and stringent manufacturing control to ensure mRNA stability and high activity in LNP delivery systems.

## INTRODUCTION

Nucleic acid-based medicines have emerged as promising alternatives to more traditional vaccines and therapeutics^1, 2^. Most notably, mRNA-based vaccines recently developed by Pfizer/BioNTech and Moderna changed the course of the SARS-CoV-2 pandemic, receiving swift emergency use authorization for public use due to efficacy and safety demonstrated in phase 3 trials ^3, 4^. The rapid deployment of these vaccines was in part due to the advantages offered by mRNA versus conventional vaccines, including flexibility of mRNA sequence design and scalability of the manufacturing process. Furthermore, the rapid biodegradability of mRNA makes it an appealing modality from a safety and pharmacokinetic perspective^5^; however, this same intrinsic instability is a key shelf-life limiting parameter and hurdle to effective vaccine delivery through various routes of administration^6^.

The effective delivery of mRNA-based vaccines and therapies is enabled by the use of lipid nanoparticles (LNPs), which protect nucleic acid degradation by exo- and endonucleases^7, 8^ and facilitate cellular uptake and expression^9, 10^. Used in both the Pfizer/BioNTech and Moderna COVID-19 mRNA vaccines, this delivery system is particularly effective as it leverages LNP surface properties^11-14^, the ability of LNPs to facilitate endosomal escape through ionization of the amino lipid^15, 16^, and deliver mRNA to specific tissues based on particle size^17^. Together, these features improve vaccine immunogenicity^18^.

Although LNP technology is an effective route for mRNA delivery to tissues, the interaction of certain chemical functionalities during storage such as oxidation, hydrolysis, or transesterification can lead to mRNA degradation^19, 20^ through backbone cleavage of the mRNA into smaller fragments^21^.

Herein we report the discovery, characterization, and identification of another class of mRNA reactivity that leads to loss in activity: the formation of lipid-mRNA adducts through the covalent addition of reactive lipid species to the nucleobase. Importantly, as many lipid-based nucleic acid formulations share common chemical functionalities, particularly those that use an ionizable amino-lipid, mechanisms identified in this study are broadly applicable. These data can inform manufacturing protocols to limit the formation of lipid-mRNA adducts and ensure the high quality of nucleic acid-based products.

## RESULTS

### Discovery of lipid-modified mRNA species in formulated mRNA-LNP system

Reversed phase–ion pair high performance liquid chromatography (RP-IP HPLC) is a well-accepted method alongside agarose or polyacrylamide gel or capillary electrophoresis (CE) to assess mRNA integrity^22, 23^. As in more conventional RP chromatography modes, separation of phases is driven by hydrophobic interactions between the analyte and stationary phase; however, since the phosphodiester mRNA backbone is highly polar, high salt concentrations are necessary to neutralize the negative charge and allow retention based on hydrophobicity of the aromatic nucleobases. Alkylammonium salts can increase hydrophobic interactions, driving selectivity based on the number of charges conferred by sequence length and enabling high resolution size-based separations (Figure 1a). This provides a similar separation to CE, which is driven by size and charge; however, based on the ion pair system used, RP-IP retains some selectivity to variations in mRNA hydrophobicity due to sequence or chemical modifications^24^.

**Figure 1.**
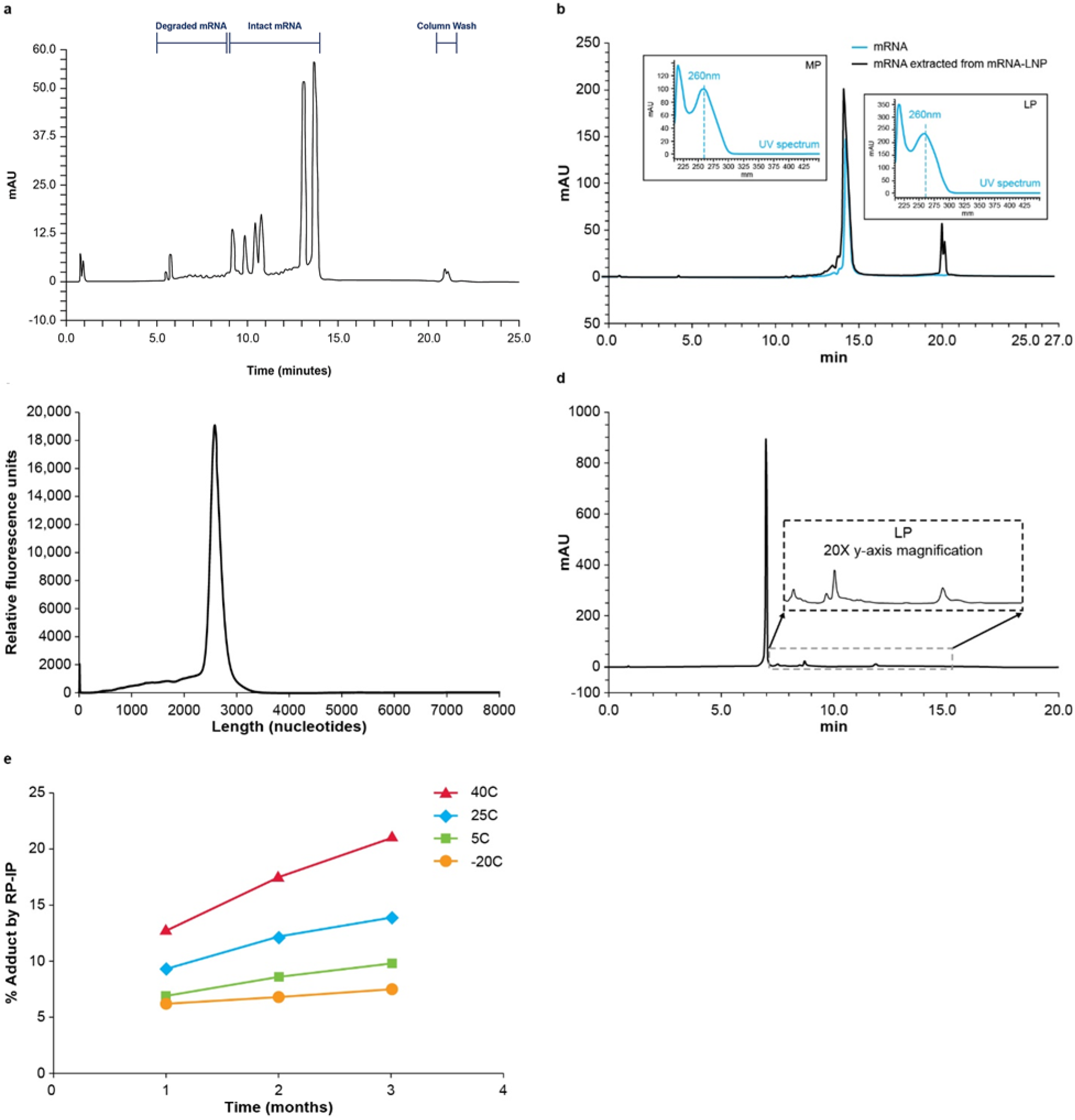
Identification of LP in formulated mRNA-LNP by RP-IP HPLC. **a**, Reversed phase ion pair high performance liquid chromatography (RP-IP HPLC) provides high resolution mRNA length-based separations to assess content and quality of mRNA products, as shown by the separation of 6 mRNAs of different lengths (659, 785, 914, 1106, 2498, and 2993 nucleotides) in a single lipid nanoparticle (LNP) formulation across retention times of 9.5–15 minutes. Peaks prior to 9.5 minutes are degradants and the peaks at 22 minutes form part of the column wash. **b**, The RP-IP HPLC analysis of pure mRNA (blue) yields a single MP (retention time 15 min), with shorter degradation products and impurities eluting prior (retention time, 10– 14.5 min), while mRNA extracted from an mRNA-LNP formulation (black) yields an additional late-eluting peak region (LP; retention time 19–21 min). The UV spectrum at each peak apex obtained from an on-line 3D UV detector shows an identical profile with maximum absorbance of 260 nm (inset). **c**, CE analysis of extracted mRNA shows a single peak, with no additional late-eluting species. **d**, LP species in mRNA extracted from an mRNA-LNP formulation can be further resolved with adjusted gradient conditions and show a polydisperse fingerprint of species. **e**, An mRNA-LNP formulation was stored for three months at four different temperatures (40°C, red; 25°C, blue; 5°C, green; -20°C, orange), and sampled at 1, 2, and 3 months for analysis by RP-IP HPLC; each data point is a single incubation condition run in a single RP-IP HPLC assay. Representative data are shown. AU, absorption units.

When RP-IP HPLC integrity analysis was applied to mRNA extracted from an mRNA-LNP, a late eluting-peak (LP) was detected by HPLC (Figure 1b) that was not observed by CE (Figure 1c). The elution time for the LP was 21 minutes, distinct from the mRNA elution time of 10– 16 minutes (Figure 1b). The ultraviolet (UV) spectrum of the LP had the same maximum absorbance at 260 nm as the mRNA main peak (MP), confirming the LP to be an RNA-related population (Figure 1b inset). Equivalent levels of the LP were observed under various analytical conditions, including modified RNA extraction, RP-IP mobile phase conditions, and stationary phase selection and temperature, suggesting it was not a separation artifact.

To further understand the late eluting time of the LP compared with various-length mRNAs, modified gradient conditions were applied, revealing the LP to be a heterogenous mixture of resolved, well defined-peaks with a characteristic fingerprint (Figure 1d). Higher levels of the LP from mRNA-LNP formulations were observed over time at higher storage temperatures (Figure 1e).

### Characterization of the modified mRNA fraction

mRNA was extracted from LNPs and fractionated by RP-IP HPLC to generate purified MP and LP fractions (Supplementary Figure 1). Upon reanalysis, separation of the MP and LP was preserved when assessed by RP-IP HPLC (Figure 2a), but fractions were not distinguished by CE (Figure 2b).

**Figure 2.**
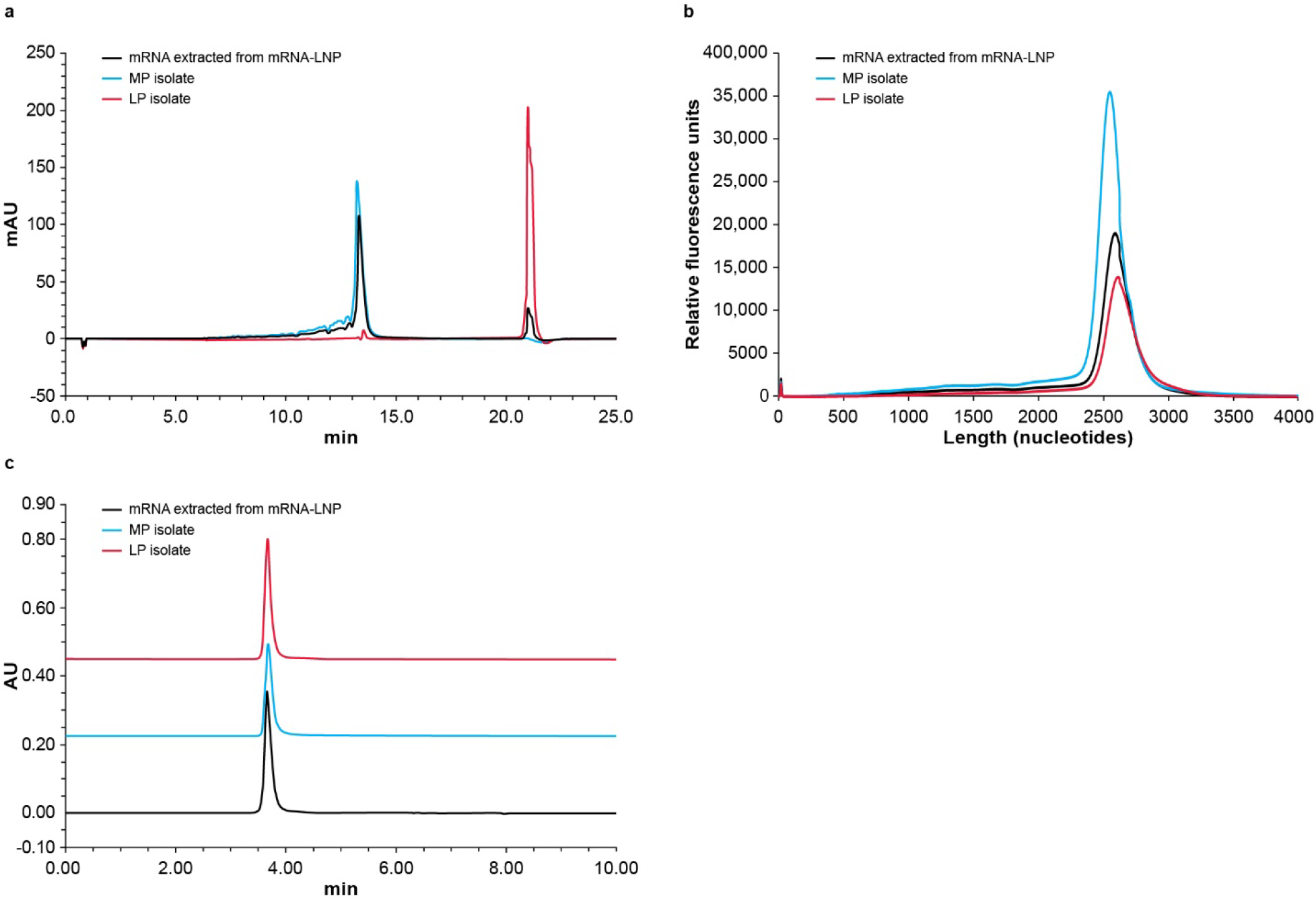
Second dimension intact analysis of isolated MP and LP. **a**, The reversed phase ion pair high performance liquid chromatography (RP-IP HPLC) chromatogram of mRNA extracted from the mRNA-lipid nanoparticle (mRNA-LNP, black) is overlaid with RP-IP HPLC re-analysis of isolated mRNA main peak (MP, blue) and late eluting-peak (LP, red), showing preserved retention time of each isolated region. **b**, The capillary electrophoresis (CE) electropherogram of mRNA extracted from the mRNA-LNP (black) is overlaid with electropherograms of the isolated MP (blue) and LP (red), showing no difference in migration time. **c**, The size-exclusion chromatography (SEC) chromatograms of mRNA extracted from the mRNA-LNP (black) is overlaid with SEC chromatograms of isolated MP (blue) and LP (red), showing no presence of aggregation in either fraction. Representative data from 4 repeat experiments are shown. AU, absorption units.

To investigate whether tertiary mRNA structures played a role, size exclusion chromatography (SEC) at ambient conditions was applied to the MP and LP. The extensive intra- and inter-molecular structure of mRNA molecules leads to a sequence-, salt-, and temperature-dependent ensemble of structures; these are typically denatured under RP-IP HPLC or CE conditions but can be resolved by native SEC^25^. SEC analysis revealed that the MP and LP fraction profiles were identical to mRNA extracted from the LNP, with a dominantly monomeric profile (Figure 2c). Together, these results eliminated aggregation as the origin of the LP and strongly implicate additional hydrophobicity behind this phenomenon.

We next applied compositional analysis to differentiate the LP from MP. The UV spectral data did not distinguish LP from MP (Figure 1b). Fourier-transform infrared spectroscopy (FT-IR) revealed minor differences potentially consistent with chemical modification (Supplementary Figure 2). Next-generation sequencing (Supplementary Figure 3) and RNA oligonucleotide mapping by digest-based mass spectrometry (MS; Supplementary Figure 4) showed an identical profile between the MP and LP, suggesting non-site-specific modifications with low abundance. Nucleotide profiling by LC-UV showed identical composition of nucleobases (Supplementary Figure 5); MS analysis did not identify LP modifications, potentially due to poor ionization of lipid modified nucleotide species in negative electrospray ionization mode.

### Identification of lipid-modified nucleosides

Nucleoside profiling was performed by enzymatic digestion of the LP and MP fractions and analysis using positive mode LC-MS/MS. Although UV and total ion-current chromatograms showed identical composition of four unmodified nucleobases for the MP and LP fractions, differential analysis revealed several mass-to-charge (m/z) values exclusively found in the isolated LP, with <1% abundance relative to total unmodified nucleosides (Figure 3a). Under LC conditions used, elution time of these unique masses was 9.5–11 minutes; this elution time was later compared with unmodified nucleosides (2–8 minutes) and earlier compared with ionizable lipid (12 minutes), indicating the unique masses had an intermediate hydrophobicity between the unmodified nucleosides and ionizable lipid.

**Figure 3.**
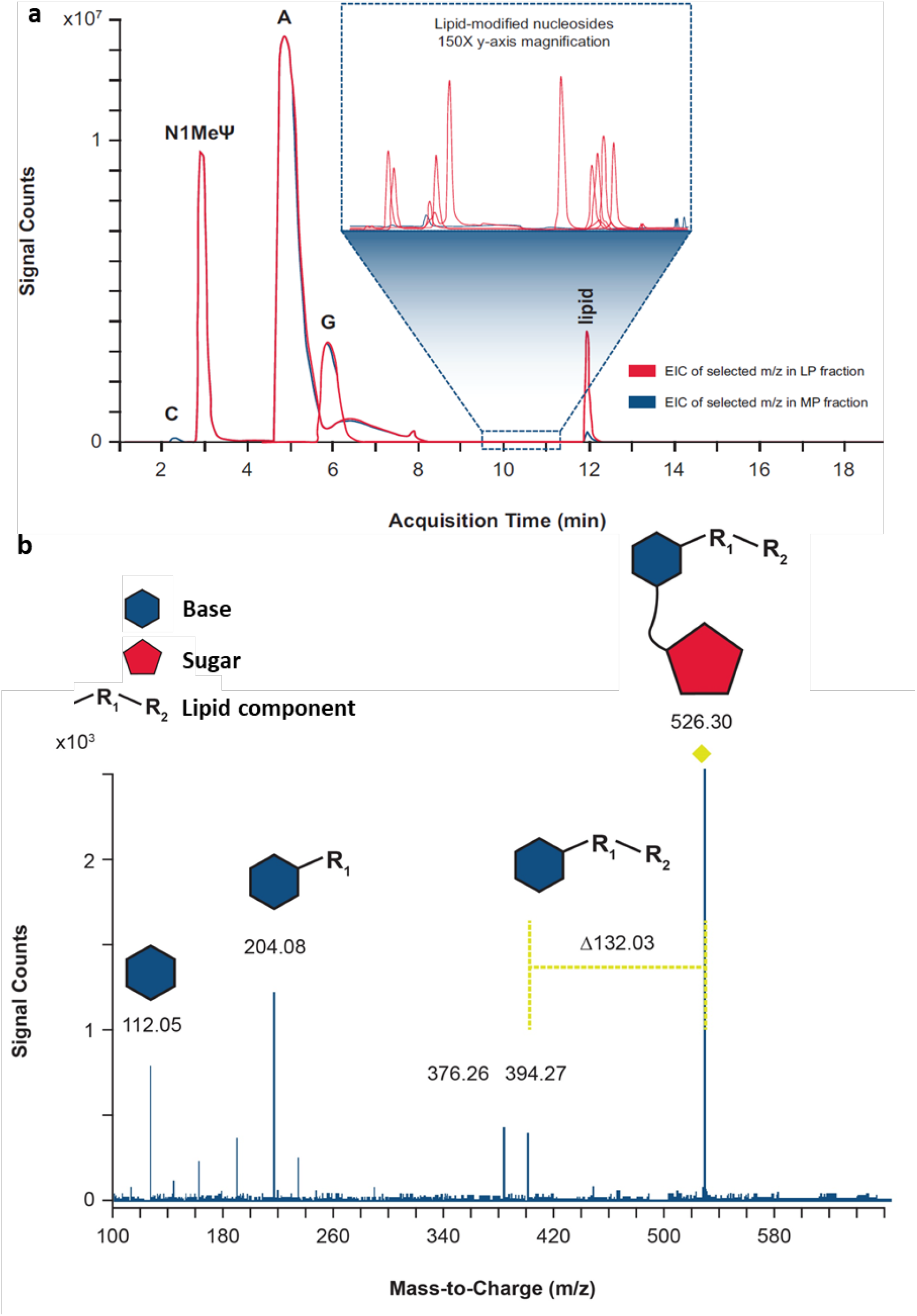
Identification of single modified nucleosides by LC/MS/MS. **a**, Isolated mRNA main peak (MP) and late eluting-peak (LP) were enzymatically digested to single nucleosides and analyzed by liquid chromatography with tandem mass spectrometry (LC/MS/MS). Extracted ion chromatograms (EICs) of selected mass-to-charge ratios (m/z) corresponding to various unmodified nucleosides, lipid-modified nucleosides, and carry-over lipid in LP (red) and MP mRNA (blue) fractions are overlaid. The selected m/z for EICs include unmodified nucleosides (Cytidine [C; 2.3 min]. N1-methylpseudouridine [N1-MelJ.I; 3 min], Adenosine [A; 5 min]. Guanosine [G; 6 min]), lipid (11.9 min), and several lipid-modified nucleosides (9.5–11.5 min) observed in LP mRNA fractions. 150x y-axis zoom of various low-abundant lipid-modified nucleosides eluting between 9.5 and 11 minutes is shown in the inset. **b**, The precursor ion at m/z 526.30 was isolated and subjected to collision-induced dissociation. Based on fragmentation pattern, the original nucleoside was determined to be cytidine, covalently modified by one chain of the ionizable lipid prepared in the binary. The fragment ion of m/z 112.05 corresponds to the exact mass of protonated cytosine (nucleobase). The characteristic neutral mass loss of 132.03 Da, corresponds to the monoisotopic residue mass of ribose.^26^ The fragment ion of m/z 394.27 corresponds to the lipid-modified cytosine, which further undergoes fragmentation to m/z 376.26 (loss of water) and 204.08 (at the internal ester), confirming the identity of the lipid chain. This cytidine modification is provided as a representative example, but similar characteristic neutral mass losses of 132.03 were observed for lipid modifications across all four nucleobases. Representative data from 4 repeat experiments are shown.

To elucidate the structure of each reacted nucleoside detected, elemental composition was assigned based on high-resolution MS data. Further MS/MS analyses across reacted nucleosides showed covalent addition of additional mass via the nucleobase, as shown by the fragmentation pattern of the ion at m/z 526.30 (Figure 3b). The presence of a fragment ion at m/z 112.05 corresponds to the protonated monoisotopic mass of cytosine. The neutral mass loss of 132.03 Da corresponds to the monoisotopic residue mass of ribose^26^, indicating that ribose is unmodified. The cytosine adduct formation serves as only one example; across several lipid systems studied, a variety of lipid adducts were observed, with reactions across all four mRNA nucleobases demonstrated in model systems (Supplementary Figure 6).

### Reaction modeling to identify contributors to adduct formation

To investigate the source of above-mentioned reactions, combinations of mRNA with individual LNP components (ionizable cationic lipid, polyethylene glycol [PEG] lipid, sterol, and phosphocholine) were examined. LP was only observed in preparations with the ionizable lipid component, with no impact from other lipids (Figure 4a). When the RP-IP HPLC chromatogram of a binary combination of mRNA and ionizable lipid was compared to that of mRNA-LNP formulated with the same ionizable lipid, an identical LP peak profile was observed by RP-IP (Figure 4b), suggesting the same reactions were occurring in this simplified system.

**Figure 4.**
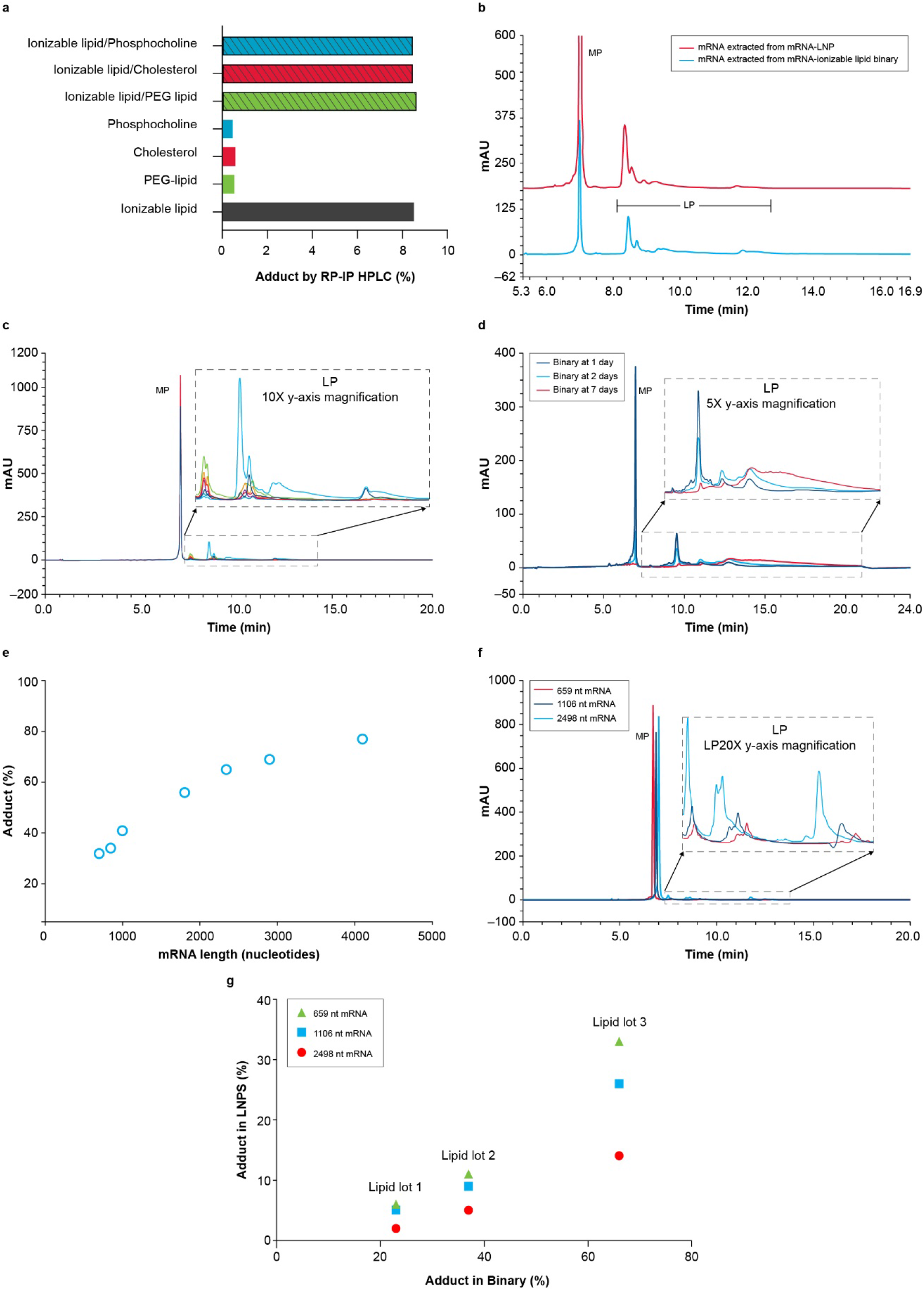
Contribution of mRNA and lipid to adduct formation. **a**, Binary reactions of single lipid components shows only combinations including the ionizable lipid resulted in significant adduct formation with mRNA by reversed phase ion pair high performance liquid chromatography (RP-IP HPLC). PEG, polyethylene glycol. **b**, The RP-IP HPLC adduct profiles of mRNA extracted from the binary system (blue) and mRNA-lipid nanoparticle (mRNA-LNP, red) with the same ionizable lipid show the same qualitative peak profile. mRNA main peak (MP). **c**, Seven different lots of the same ionizable lipid were prepared in binaries with mRNA, yielding a variety of peak profiles and abundances of adduct species in the overlaid RP-IP HPLC chromatograms. **d**, Adduct formation in a binary reaction with a highly reactive lipid was evaluated by RP-IP HPLC at 1 day (black), 2 days (blue), and 7 days (red), showing increased tailing likely corresponding to the accumulation of multiple adducts per mRNA molecule. **e**, Adduct formation in binary reactions was evaluated with mRNA molecules of different lengths by RP-IP HPLC at equivalent mRNA masses. An increase in late eluting-peak (LP) with length is consistent with a constant rate of modification on the single nucleotide level. **f**, RP-IP HPLC chromatographs of 659 (red), 1106 (dark blue), and 2498-nucleotide (blue) mRNAs show an increase and left shift of each adduct peak with increasing mRNA length. **g**, Adduct formation as a function of mRNA length was assessed by RP-IP HPLC for mRNA-LNP and corresponding binaries. A positive correlation was observed, with more adduct at longer mRNA lengths. Representative data from ≥3 repeat experiments are shown. AU, absorbance units.

Additional ionizable lipid chemistries (>100) were next examined, all of which generated quantifiable levels of mRNA-lipid adduct, demonstrating a broad class effect. The RP-IP HPLC profile of these adduct species varied in retention times across different chemistries likely based on size and hydrophobicity of the adducted lipid structure (Supplementary Figure 7). Across different lots of ionizable lipid tested in the binary system, there were variations in relevant abundance of a consistent set of LP species (Figure 4c), suggesting these reactions are driven by impurities of the ionizable lipid rather than chemical reactivity of the lipid itself. Subsequently, a highly reactive ionizable lipid from the chemistry screen was studied to understand the chromatographic behavior of intact adducted mRNA. At 1 day, a discrete shift in retention produced LP peaks at 10–12 minutes; increased tailing from 12.5–22.5 minutes was observed as the MP was depleted (Figure 4d). This observation strongly suggests a single adduct event can shift the RNA molecule to LP, and accumulation of multiple adducts per mRNA molecule further drives increases in retention time.

The binary system was then used to study contributions of the mRNA molecule to adduct formation. A series of mRNA sequences ranging in length from 700 to >4000 nucleotides in length was assessed at equivalent masses in individual binary reactions, resulting in increasing LP with mRNA length (Figure 4e). As relative UV intensity of the LP peak in the RP-IP HPLC assay is correlated with relative mass, this observation is consistent with a constant rate of reaction on the single base level resulting in greater mass shifts to LP with increasing sequence length. This correlation highlights the relevance of these reactions for mRNA as a high molecular weight nucleic acid polymer. Although comparable lipid systems are used in low molecular weight RNA products such as small interfering RNA (siRNA), the same molar reaction rate observed in mRNA systems would result in lower levels of adduct formation of siRNA on an intact mass basis due to the stochastic nature of adduct formation. Notably, the same adduct peaks can be observed across mRNAs of different lengths in the chromatographic profile with a decrease in retention time of each LP region with increasing mRNA length (Figure 4f). We can understand this as an average hydrophobicity of the adducted mRNA sequence; as sequence length increases, the same adducts comprise less of the total molecule, lowering the impact on retention. The combined impact of mRNA length and lipid lot is shown in Figure 4g, demonstrating that the simplified binary system can be used as a predictive model for the final LNP.

### Oxidative impurities of ionizable lipid as a driver of adduct formation

To further understand the implicated role of ionizable lipid impurities as the driver of adduct formation (Figures 4a-c), impurities and degradants that are common to the ionizable amino-lipid family such as those shown in Figure 5a were next examined. N-Oxide formation is a common degradation pathway for tertiary -amine -containing molecules under oxidative stress^27^. Although relatively stable, N-oxide can further hydrolyze to secondary amines and aldehyde counterparts (Figure 5b), reported typically through metal catalysis^28^. Since transition metals are well-controlled in mRNA-LNP raw materials due to their propensity to accelerate mRNA degradation, the relevance of N-oxide hydrolysis in the absence of metal catalysts was evaluated. N-oxide standard was generated from a representative ionizable lipid, precipitated in acidic buffer, and analyzed by reversed phase ultra-high-performance liquid chromatography with charged aerosol detection (RP-UPLC-CAD) and MS/MS detection. Three peaks generated under these conditions at retention time 8.7, 10.3, and 25 minutes were identified by MS/MS as the three secondary amines resulting from hydrolysis of N-oxide. To detect the residual mass, the degraded N-oxide solution was derivatized with an aminooxy-PEG to label all aldehyde functionalities. Unique peaks at 9.5 and 16 minutes were detected by RP-UPLC-CAD, corresponding to aldehyde products of N-oxide hydrolysis; the third expected aldehyde peak likely elutes in the column void (Figure 5c). These studies demonstrate that even in relatively mild acidic conditions, N-oxide hydrolysis can generate secondary amines and aldehydes without the use of metallic catalysts. Further analysis revealed the presence of N-oxide leads to high levels of LP, with almost complete conversion of the mRNA to LP within 3 days in the binary system (Figure 5d).

**Figure 5.**
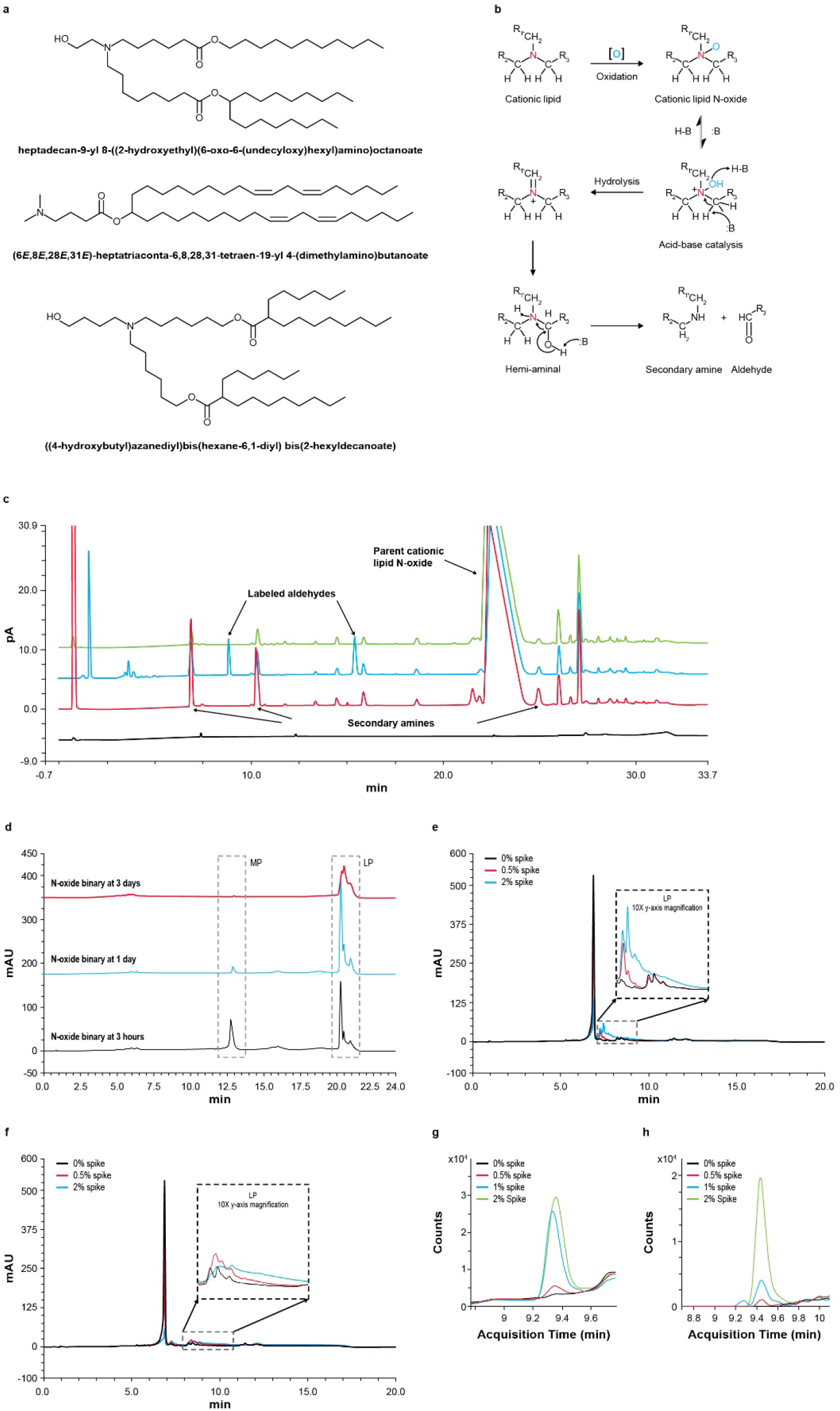
N-oxide as a driver of adduct formation. **a**, Tertiary-amine-containing ionizable lipids currently used in RNA-lipid nanoparticle (RNA-LNP) products. **b**, N-oxide forms through tertiary amine oxidation and undergoes an acid/base-catalyzed hydrolysis at the amine to generate aldehydes and secondary amines. **c**, N-oxide acid hydrolysis products were detected by reversed phase ultra-high-performance liquid chromatography with charged aerosol detection and tandem mass spectrometry (RP-UPLC-CAD MS/MS); N-oxide standard (green), acid-precipitated N-oxide standard with (blue) and without (red) aminooxy-polyethylene glycol (aminooxy-PEG) label, and buffer baseline (black). Secondary amines from N-oxide hydrolysis elute at 8.7, 10.3, and 25 minutes. Corresponding aminooxy-PEG-derivatized aldehydes elute at 9.5 and 16 minutes, with the third likely in the column void. pA, pico ampere. **d**, Binary reactions of mRNA with pure N-oxide result in high late eluting-peak (LP). RP-IP analysis of mRNA extracted from the binary is shown at 3 hours (black), 1 day (blue), and 3 days (red) MP, mRNA main peak. **e and f**, Binary reactions spiked with (d) pure 17-carbon aldehyde and (e) pure 25-carbon aldehyde products from the N-oxide degradation pathway result in high levels of mRNA modification. Reversed phase ion pair high performance liquid chromatography (RP-IP HPLC) chromatograms of the extracted mRNA are shown with no aldehyde spike (black), 0.5% aldehyde (red), and 2% aldehyde (green). **g and h**, Binary reactions with the pure 17-carbon aldehyde from the N-oxide degradation pathway were analyzed for single nucleoside modifications. Binaries were prepared with no aldehyde (black), 0.5% aldehyde (red), 1% aldehyde (blue), and 2% aldehyde (green) spike. mRNA was extracted, enzyme-digested, and analyzed by liquid chromatography with tandem mass spectrometry (LC-MS/MS). The mass-to-charge ratios (m/z) corresponding to aldehyde-cytidine adducts (m/z 526.3 and 540.3) increased with aldehyde spike level. A representative ionizable lipid system, heptadecan-9-yl 8-((2-hydroxyethyl)(6-oxo-6-(undecyloxy)hexyl)amino)octanoate is used here. Representative data from ≥3 repeat experiments are shown.

### Investigation of mRNA adduction by aldehydes

Covalent modification of DNA by aldehydes has been previously reported,^29^ setting a precedent for the reactivity of newly identified aldehydes from N-oxide degradation of amino lipids. Two representative aldehydes from the N-oxide degradation pathway of an ionizable lipid (a linear, 17-carbon chain and a branched, 25-carbon chain) were studied. The aldehydes were individually spiked into ionizable lipids as impurities prior to precipitation in binary reactions. Significant increases in lipid-mRNA adduct levels were observed with increasing spike concentration for both aldehydes. By RP-IP HPLC, peaks corresponding to mRNA adducts with the smaller linear aldehyde elute at 7.5 minutes (Figure 5e) and those for the larger branched aldehyde elute at 8–10 minutes (Figure 5f). When mRNA from the linear aldehyde binary was subjected to enzymatic digestion and positive mode LC-MS/MS analysis, there was a corresponding increase in 2 resulting nucleoside masses of m/z 526.3 (Figure 5g) and 540.3 (Figure 5h) which were further elucidated by MS/MS as being products of aldehyde addition to cytidine (Figure 3b). As described above (Figure 4d), the increase in tailing at the highest spike concentration in the intact mRNA chromatography is potentially due to the accumulation of multiple lipid adducts per mRNA strand, contrasting the initial discrete shift that occurs for each aldehyde with a single adduct addition. Although hydrolysis of N-oxide is one particularly relevant pathway for the generation of aldehydes in the LNP system, similar species may be present as lipid raw material impurities or oxidative degradants. These could generate the diversity of adduct species observed within the intact mRNA and single nucleoside level.

### Impact of lipid-mRNA adduct on protein expression and product shelf life

The role of chemical modifications on protein expression was explored. MP and LP fractions were isolated by RP-IP HPLC from 2 mRNA formulations encoding the human erythropoietin (hEPO) protein and analyzed for *in vitro* protein expression in BJ fibroblasts. Expression of hEPO protein was comparable between the isolated MP, the unfractionated extracted mRNA, and hEPO assay control (Figure 6a); in contrast, no protein expression was detected from the isolated LP (Figure 6a). Separately, 5 LNP formulations were subjected to various levels of degradation, after which the mRNA was extracted from the LNP and analyzed by RP-IP HPLC, CE, and for *in vitro* protein expression in HeLa cells (Figure 6b). When mRNA integrity was assessed using RP-IP HPLC, a correlation between relative mRNA integrity (including the adduct as an impurity) and protein expression was found (Figure 6b). In contrast, mRNA integrity measured by CE only weakly correlated with protein expression; all samples had a narrow range of relative purity despite the decrease in protein expression (Figure 6b). Extrapolating these correlations show a total loss in protein expression despite mRNA purity >60% by CE. These data demonstrate the potential of such adduct reactions to reduce activity of mRNA-LNP products and the inadequacy of CE to determine mRNA quality in LNP formulations.

**Figure 6.**
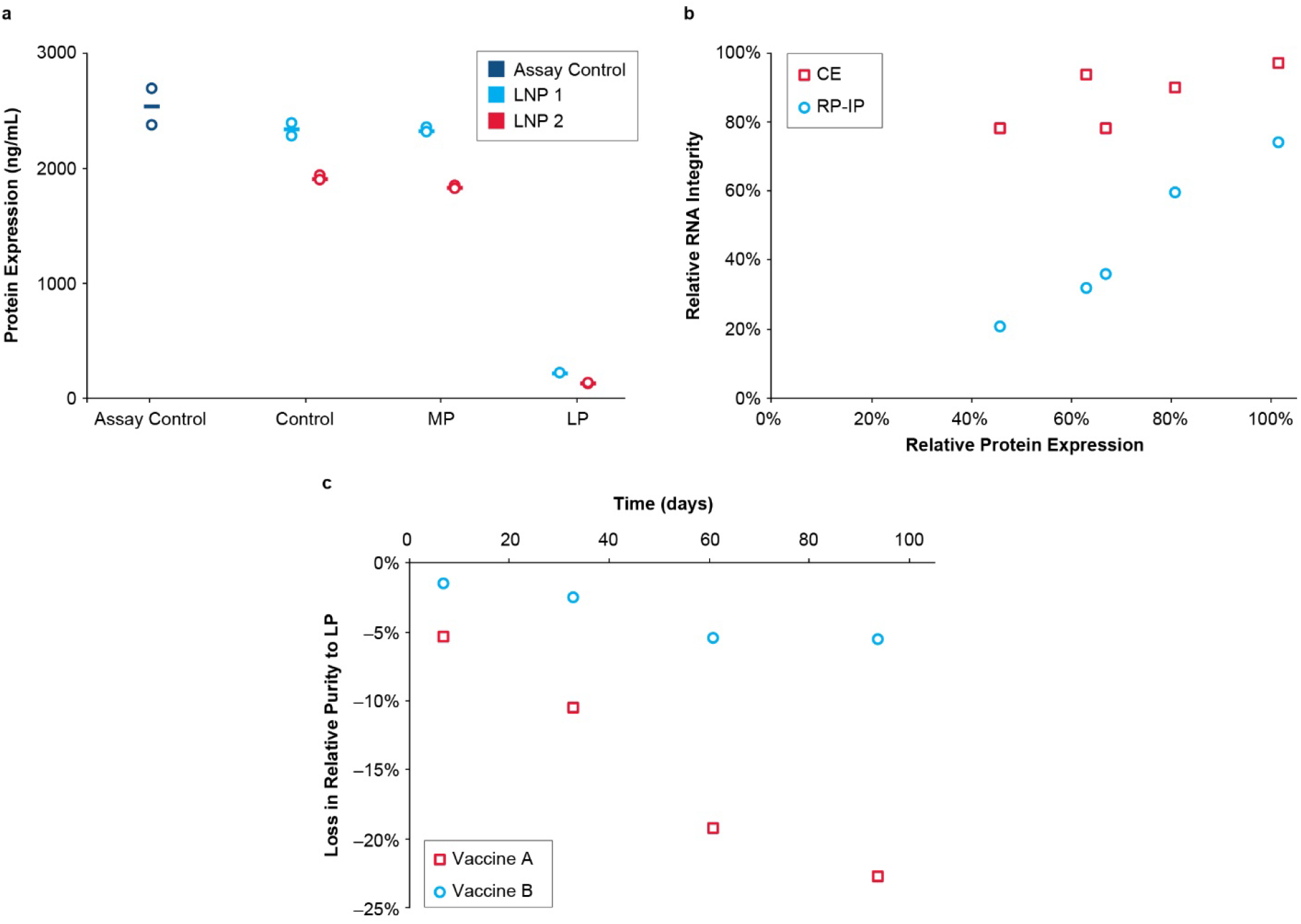
mRNA adduction reduces protein expression. **a**, Isolated main peak (MP) and late eluting-peak (LP) alongside control mRNA-lipid nanoparticle (mRNA-LNP) and an assay positive control were tested for *in vitro* protein expression in BJ fibroblasts after 48 hours. RNA was extracted from 2 hEPO-LNP formulations by isopropanol precipitation and purified by RP-IP to generate MP and LP prior to transfection. The assay control is a pure hEPO mRNA standard, the control sample is mRNA extracted from each formulation prior to RP-IP separation, and the MP and LP are isolated fractions. Results from 2 replicate transfection wells is plotted with the mean. **b**, Five different mRNA-LNP samples were prepared using different ionizable lipids and incubated at 5°C to generate varying levels of adduct and degradation. RNA was extracted from the mRNA-LNP by isopropanol precipitation and evaluated by reversed phase-ion pair (RP-IP), capillary electrophoresis (CE), and *in vitro* protein expression in HeLa cells as measured by mean fluorescence intensity. Relative expression as a percentage of the neat mRNA expression is plotted versus relative integrity as a percentage of the neat mRNA integrity, determined by relative area using both CE and RP-IP HPLC. **c**, Loss in mRNA purity to adduct formation in two vaccine formulations is plotted over 3 months at refrigerated conditions. Poor process control led to high LP in Vaccine A, but adduct was well-controlled in Vaccine B. Representative data are shown for 3 repeat experiments for Figure 6a and multiple repeat experiments for Figures 6 b and c.

The introduction of impurities into the vaccine manufacturing process could potentially impact vaccine stability even under refrigerated conditions. The loss of mRNA purity to adduct formation under refrigeration (5°C) was examined for 2 vaccine formulations utilizing the same mRNA sequence and lipid system. One vaccine had poor controls in place to limit adduct formation (Vaccine A) and the other had rigid controls (Vaccine B; Figure 6c). Data for Vaccine A show an initial delta of 15%, indicating rapid adduct formation during product processing or prior to testing. This is followed by substantial loss in mRNA integrity by almost 50% to adduct formation over 3 months in refrigeration. By contrast, Vaccine B demonstrated comparatively low levels of adduct formation with a negligible increase over time (Figure 6c).

## DISCUSSION

LNP-based nucleic acid delivery systems enabled the development and emergency use authorization of 2 COVID-19 mRNA vaccines at unprecedented speed. The agility offered by the mRNA-LNP platform promises a new era in vaccine and drug development. Ionizable lipids are an essential component of the LNP; they aid in encapsulating mRNA into the LNP system, define the biophysical profile and stability, and determine pharmacological performance of the vaccine or therapeutic.

Herein we report identification of a lipid-modified class of mRNA impurities generated by electrophilic degradants and impurities originating in the ionizable lipid; these impurities disrupt mRNA translation and negatively impact the activity of LNP-formulated mRNA products. Additionally, such impurities are difficult to identify using traditional techniques used to assess mRNA integrity in mRNA-LNP such as CE. RP-IP HPLC provides specificity and sensitivity to detect adducted mRNA molecules, which are otherwise difficult to identify due to the low rate of modification. Indeed, data suggest that RP-IP HPLC can detect even single adduct events on intact mRNA, whereas in contrast, even purified LP is not distinguished by the conventional CE methodology. Furthermore, it is critical that formation of these mRNA-lipid adducts are examined during formulation design, clinical evaluation, and commercialization by using appropriate RP-IP HPLC methodologies to ensure there is no loss in mRNA integrity.

A potential source of impurities is through hydrolysis of N–Oxide to aldehydes; this reaction is broadly relevant to tertiary amines commonly used in LNP formulation of siRNA and mRNA^28, 30^. It remains highly probable that formation of this class of adducts has been missed by historically applied analytical technologies such as CE, thus overlooking an important critical quality attribute of mRNA-LNP. This represents a gap in quality control of mRNA-LNPs during manufacturing, particularly as it pertains to consistency and activity of the resultant drug product. In this study, we report the ability of RP-IP HPLC to resolve aldehyde species differing only by a short alkyl chain, highlighting the selectivity of the RP-IP HPLC method to detect minute differences on the intact mRNA molecule.

The pathway reported herein identifies an important class of impurities that may be present in all lipid-based mRNA systems. These reactions can be mitigated through raw material control, manufacturing process parameters, formulation design, and LNP storage conditions. It thus remains critical to monitor and control lipid adduct formation during the research, development, and manufacture of LNP-formulated nucleic acid products. Importantly, this study highlights the need for advanced methods such as RP-IP HPLC to ensure the quality, consistency, and efficacy of any pharmaceutical product.

## METHODS AND MATERIALS

### Formulations

All mRNA and LNP formulations used in this work were representative of GMP-grade material. Several mRNA molecules were used throughout this work; all were 5-prime capped, 3-prime poly-adenylated sequences with modified uridine chemistry (N1-methyl-pseudouridine) encoding different vaccine targets and other proteins.

### RNA extraction from mRNA-LNPs and binaries

mRNA was extracted from the mRNA-LNP formulation or lipid binary mixture by isopropanol precipitation. 100 µL of mRNA-LNP or binary was diluted 10-fold in 900 µL ammonium acetate (60 mM) in isopropanol, vortexed briefly, and centrifuged at 14,000g for 15 minutes at 4 °C. The supernatant was discarded and the pellet was washed with 1 mL isopropanol, vortexed, and centrifuged at 4 °C; the pellet was dried in vacuo and resuspended in 100 µL RNase-free water at room temperature.

### Reversed phase ion pair chromatography (RP-IP)

Sample separation was performed on a DNAPac RP column with 4-µm particles and dimensions of 2.1 × 100 mm (Thermo Fisher Scientific) at a flow rate of 0.35 mL/minute and column temperature of 65 °C. Mobile phase A consisted of dibutylammonium acetate (50 mM; TCI America) and triethylammonium acetate (100 mM; Sigma-Aldrich) and mobile phase B consisted of 50% acetonitrile (Sigma-Aldrich), dibutylammonium acetate (50 mM), and triethylammonium (100 mM). Separation was accomplished by step-gradient with an initial 1.5-minute hold at 25% B, a 3.0-minute gradient from 25-50% B, a 14.5-minute gradient from 50-56% B, and a 0.5-minute gradient and hold at 100% B. Modified gradient conditions for LP resolution were performed with an initial 1.5-minute hold at 25% B, a 1-minute gradient from 25-45% B, a 12.5-minute gradient from 45-100% B, and a 0.5-minute gradient and hold at 100% B. Approximately 2 µg of mRNA were injected. mRNA was detected by UV at 260 nm. LP is quantified as the relative percent of the total chromatographic peak area.

### Capillary electrophoresis (CE)

Sample separation was performed on a Fragment Analyzer (Agilent Technologies), an automated multiplexed CE system equipped with an LED light source and charged-coupled device detector. RNA was quantitatively and qualitatively analyzed using the RNA Analysis Kit (Agilent Technologies DNF-489-0500). The RNA separation gel was mixed with an intercalating dye (AATI) at a v/v ratio of 10,000:1 for use as the separation matrix. RNA was denatured at 70 °C for 2 minutes and cooled on ice prior to analysis. Denatured RNA samples were electrokinetically injected at 5 kV for 6 seconds, and electrophoresis was performed for 40 minutes at 8 kV. An RNA ladder (AATI) was similarly analyzed as a calibrator for nucleotide size. Data were analyzed using PROSize 2.0 software (Agilent Technologies).

### Size exclusion chromatography (SEC)

Sample separation was performed on a Zenix SEC-300 150 × 4.6 mm protein SEC column (Sepax) on a Waters H-Class UPLC (Waters). The mobile phase condition was 100 mM Tris acetate/2.5 mM EDTA pH 8 with an isocratic flow of 0.25 mL/minute and UV detection at 260 nm.

### Binary model preparation

mRNA-lipid binaries were formed by mixture of mRNA and ionizable lipid. Unless otherwise noted, a standard 2000-nucleotide mRNA (0.135 mg/mL) was prepared sodium acetate (37.5 mM; pH 5.3; Sigma Aldrich) and mixed at a 3:1 ratio with an ionizable lipid solution (4 mg/mL) in ethanol, followed by incubation at room temperature for 24 hours. RNA was extracted from the binary using isopropanol precipitation as described above prior to further analysis.

### Enzymatic digestion of mRNA to ribonucleosides

Total nuclease digestion was performed as previously described^31^ with modifications. mRNA was incubated with 15 units of benzonase (Millipore), 2 units of phosphodiesterase I (Sigma-Aldrich), and 1.3 units of quick calf intestinal alkaline phosphatase (New England Biolabs) in buffer containing Tris-HCl (Invitrogen), NaCl, and MgCl_2_ (Invitrogen) at 37 °C for 2 hours.

### Positive mode liquid chromatography mass spectrometry (LC-MS/MS)

Nucleosides were separated in an increasing water/acetonitrile gradient containing 0.1% formic acid at 0.4 mL/minute on an Accucore C30 column with 2.6 µm particles and dimensions of 2.1 × 250 mm (Thermo Scientific) at 50 °C. Ultraviolet detection of nucleosides was monitored at 260 nM. Mass spectral data were acquired on an Agilent 6530 QTOF (Agilent Technologies) in positive electrospray ionization (ESI) mode. The mass range was 100-2000 m/z, drying gas temperature was 290 °C at a flow rate of 11 L/min, and the nebulizer gas pressure was 35 psi. The capillary, fragmentor, and octupole voltage were set at 3500, 100, and 750 V, respectively. External calibration was used for accurate mass measurement. For tandem mass spectrometry (MS/MS), precursor ions were subjected to collision-induced dissociation and MS/MS fragmentation analysis.

### Tandem mass spectrometry (MS/MS)

MS/MS experiments using collision induced dissociation (CID) were performed following RP-HPLC separation. Acquisition of a full mass scan was followed by targeted MS/MS scans of precursor ions of interest. Data were acquired on an Agilent 6530 QTOF (Agilent Technologies) for MS/MS analysis. Normalized collision energy for ion activation was 30 arbitrary units. Narrow ion isolation width (∼1.3 m/z) was used for isolation.

### N-oxide precipitation and labeling

An N-oxide standard of ionizable lipid was generated. The N-oxide compound was dissolved in ethanol at 4 mg/mL. One milliliter of the N-oxide solution was added dropwise to 3 mL of sodium acetate (37.5 mM; pH 5.3; Sigma Aldrich), followed by incubation at room temperature for 24 hours. Prior to LC-CAD-MS analysis, aminooxy-PEG (1 mM; Thermo Fisher) was used as a labeling agent to detect and identify aldehydes present in the system.

### Aldehyde spike preparation

Synthetic aldehyde standards corresponding to the 17-carbon linear chain and 25-carbon branched chain of a representative ionizable lipid (heptadecan-9-yl 8-((2-hydroxyethyl)(6-oxo-6-(undecyloxy)hexyl)amino)octanoate) were generated. Aldehyde solutions (4 mg/mL) were individually prepared in ethanol and mixed with an ionizable lipid solution (4 mg/mL) in ethanol to various target compositions, listed in % w/w of the aldehyde to ionizable lipid. Binary preparations, intact mRNA HPLC analyses, and digested nucleoside LC/MS analyses were then performed as described above.

### Reversed phase ultra-performance liquid chromatography with charged aerosol detection (RP-UPLC-CAD)

LNP formulations were diluted in ethanol and the supernatant was analyzed. Lipid standards were separately diluted and analyzed to identify and quantify lipid components. RP-UPLC separation was performed on an ACE Excel 2 Super C18 column (Advanced Chromatography Technologies) with 2.1 × 150 mm dimensions heated to 60 °C. Mobile phase A consisted of 0.1% trifluoroacetic acid (TFA) (Thermo Fisher Scientific) in water and mobile phase B consisted of 60/40/0.1% isopropyl alcohol/tetrahydrofuran/TFA (Thermo Fisher Scientific). Lipids were eluted by step-gradient with an initial 1.5-minute hold at 5% B, a 4.5-minute gradient from 5–48% B and 4-minute hold, a 1-minute gradient from 48–56% B and 12-minute hold, and an 8-minute gradient from 56–96% B and 2-minute hold. Lipids were detected by CAD using an evaporator temperature of 35 °C and analytical gas regulation mode.

### Protein expression in BJ fibroblasts

BJ fibroblasts (ATCC; catalog #CRL-2522) were cultured in complete media (Eagle’s minimum essential medium [EMEM] with L-glutamine; 10% FBS). Cells were seeded in 96-well plates (20,000 cells/well) 24 hours prior to transfection; extracted mRNA was transfected with Lipofectamine L2000 (Life Technologies, catalog #11668019) at 250 ng/well and incubated for 48 hours at 37 °C before collection of the supernatant for analysis. A hEPO mRNA standard was included as an assay control.

### Detection of protein expression by flow cytometry

*In vitro* expression was performed in HeLa cells (ATCC, catalog #CCL-2). Cells were seeded in Minimal essential medium (MEM) with 10% FBS, 1X Glutamax, and 1X sodium pyruvate at 37 °C at 15,000 cells in a total volume of 100 µL per well and grown for 24 hours. Extracted mRNA was transfected with lipofectamine L2000 (Life Technologies, catalog #11668019) at 1333 ng of mRNA per well and incubated for 18-20 hours at 37 °C. To detect intracellular protein expression, cells were first fixed and permeabilized (Cytoxfix/Cytoperm; BD Biosciences catalog #554714) and then stained with a proprietary primary antibody (1/2000 dilution of 1 mg/mL stock). Data were acquired on the BD LSRFortessa™ Flow Cytometer (BD Biosciences) using BD FACSDiva™ version 8.0 software (BD Biosciences) and analyzed using FlowJo software version 10.4 (BD Biosciences). Mean fluorescence intensity was normalized using a neat mRNA positive control and non-translating negative control. The gating strategy can be found in Supplementary Figure 8.

### Statistics and Reproducibility

Assay measurements were taken from distinct samples. This is a descriptive study and as such no formal statistical analyses were performed. Experiments conducted for the identification and characterization of the MP and LP utilizing RP-IP HPLC, CE analysis of adduct-containing mRNA, resolution of the adduct peaks with UV spectral data, and increases in adduct at elevated storage temperature of the LNP (Figure 1) acts as a representative example of analyses that have been performed repeatedly (>100 times) with consistent reproducible findings. Experiments conducted for the isolation of adducts and analysis utilizing RP-IP, SEC and CE (Figure 2) were replicated 4 times with similar results. Identification of modified nucleosides present in the adduct by nucleoside digest, differential analysis, and subsequent structural elucidation by MS/MS (Figure 3) was replicated 4 times with similar results. For identifying the contribution of mRNA and lipids to adduct formation utilizing binary analysis of individual lipids, comparisons of ionizable lipid binary adduct results with the LNP, and qualitative observations in the binary with mRNA length and lipid lot (Figure 4) have each been replicated ≥3 times with similar results. Experiments conducted to determine the influence of N-oxides in adduct formation utilizing the decomposition of N-oxide in acidic buffer, mRNA binaries with N-oxide, and RP-IP and MS analysis of aldehyde-spiked binaries (Figure 5) have been replicated ≥3 times with similar results. Experiments to determine the expression of the isolated adduct (Figure 6a) have been replicated 3 times and experiments to determine loss in expression with adduct, increase in adduct over storage across formulations (Figures 6b and 6c) have been replicated ≥3 times.

For data shown in the supplemental information, FTIR analysis (Supplement Figure 2) was performed once. Illumina NGS (Supplemental Figure 3), T1 digest followed by LC/UV/MS (Supplemental Figure 4), and NaOH digest followed by RP-IP LC-UV (Supplemental Figure 5) were replicated 2 times with similar results. Analysis of homopolymers in the binary system (Supplement Figure 6) was performed once. Comparisons of adduct profiles (Supplement Figure 7) have been generated with >100 lipid chemistries with similar results; seven representative chromatograms are displayed.

## DATA AVAILABILITY

The authors declare that the data supporting the findings of this study are available within this Article and its Supplementary Information. The data that support the findings of this study are available from the corresponding authors upon reasonable request.

## ACKNOWLEDGEMENTS

We thank Melissa Moore, Orn Almarsson, Don Parsons, Huijuan Li, Shaila Jayaram, and Will Issa for their intellectual input and support, and Ed Miracco for assistance with the preparation of this manuscript. Jack Kramarczyk and Beth Lally led formulation efforts to apply findings to the LNP. Amelie Dricot-Ziter and Kamaljeet Sandhu supported *in vitro* expression assays. Dave Mauger performed next-generation sequencing analysis, and Christopher Tunkey and Christopher Pepin assisted with preparation of the next-generation sequencing data. Tao Jiang, Srujan Gandham, and Serenus Hua provided support on LC/MS analyses, nucleoside digestion, and differential analysis approaches and Greg Mercer and Jin Lim provided N-oxide and aldehyde standards. Medical writing and editorial assistance was provided by Wynand van Losenoord and Srividya Ramachandran, PhD, of MEDiSTRAVA in accordance with Good Publication Practice (GPP3) guidelines, funded by Moderna Inc, and under the direction of the authors.

## AUTHOR CONTRIBUTIONS

Meredith Packer and Dipendra Gyawali contributed equally to this manuscript. MP identified covalent lipid addition and developed and performed RP-IP and other intact mRNA analyses, preparative mRNA fractionation, and binary studies. DG identified the N-oxide pathway and performed binary studies, lipid analyses, and N-oxide and aldehyde analysis. RY designed and carried out LC/MS and MS/MS experiments for modified nucleoside structural analysis. JS made initial observation on the modified mRNA and provided technical expertise to N-oxide work. PW provided leadership and direction throughout the project.

