## Supplementary information for "A novel mechanism for the loss of mRNA activity in lipid nanoparticle delivery systems"

Title:

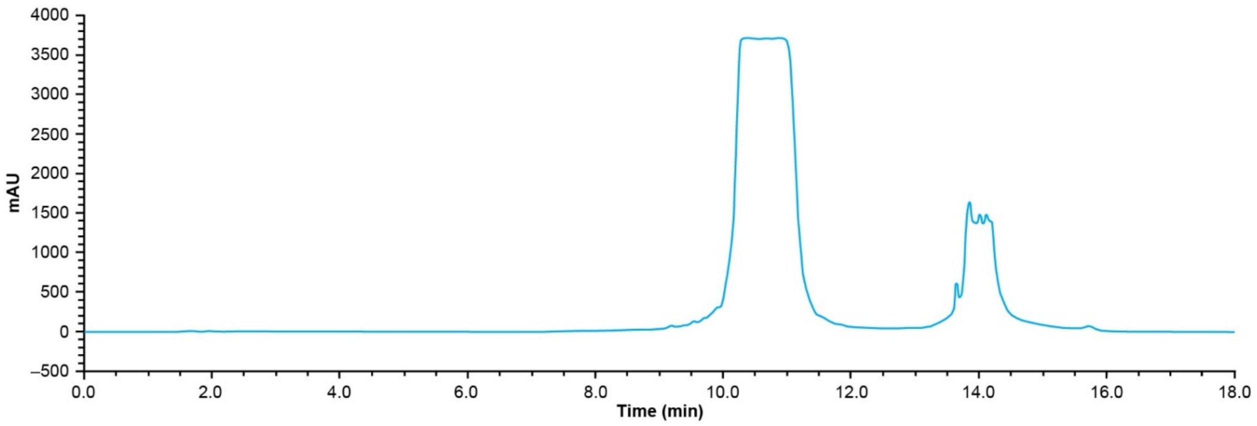


**Supplementary Figure 1.** RP-IP UV chromatogram shows the preparative separation performed to isolate MP and LP fractions with small modifications to the gradient conditions. Concentrated samples were adjusted to mobile phase A ion pair conditions, and 0.5-mL injections were performed. Fractions were collected between 10 and 12 minutes (main peak, MP) and 13.5 and 15 minutes (late eluting-peak, LP).

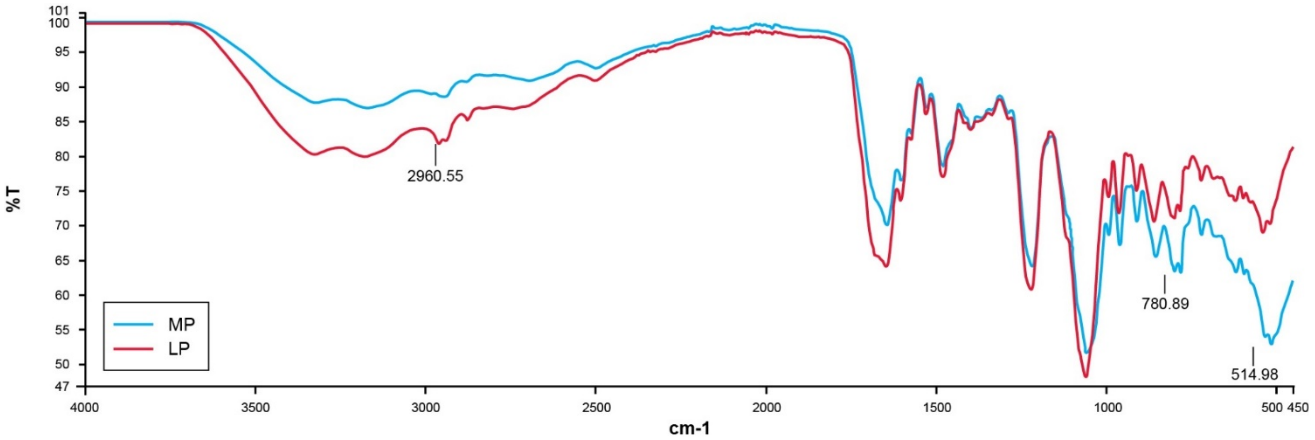


**Supplementary Figure 2.** 4 mg each of isolated main peak (MP, blue) and late eluting peak (LP, red) were evaporated to dryness in a Genevac, and the resulting film was analyzed by fourier-transform infrared spectroscopy (FTIR; n=1). Raw spectra showed similar chemical signatures with a few areas showing changes (2960.55 and the fingerprint region 780.89 - 514.98).

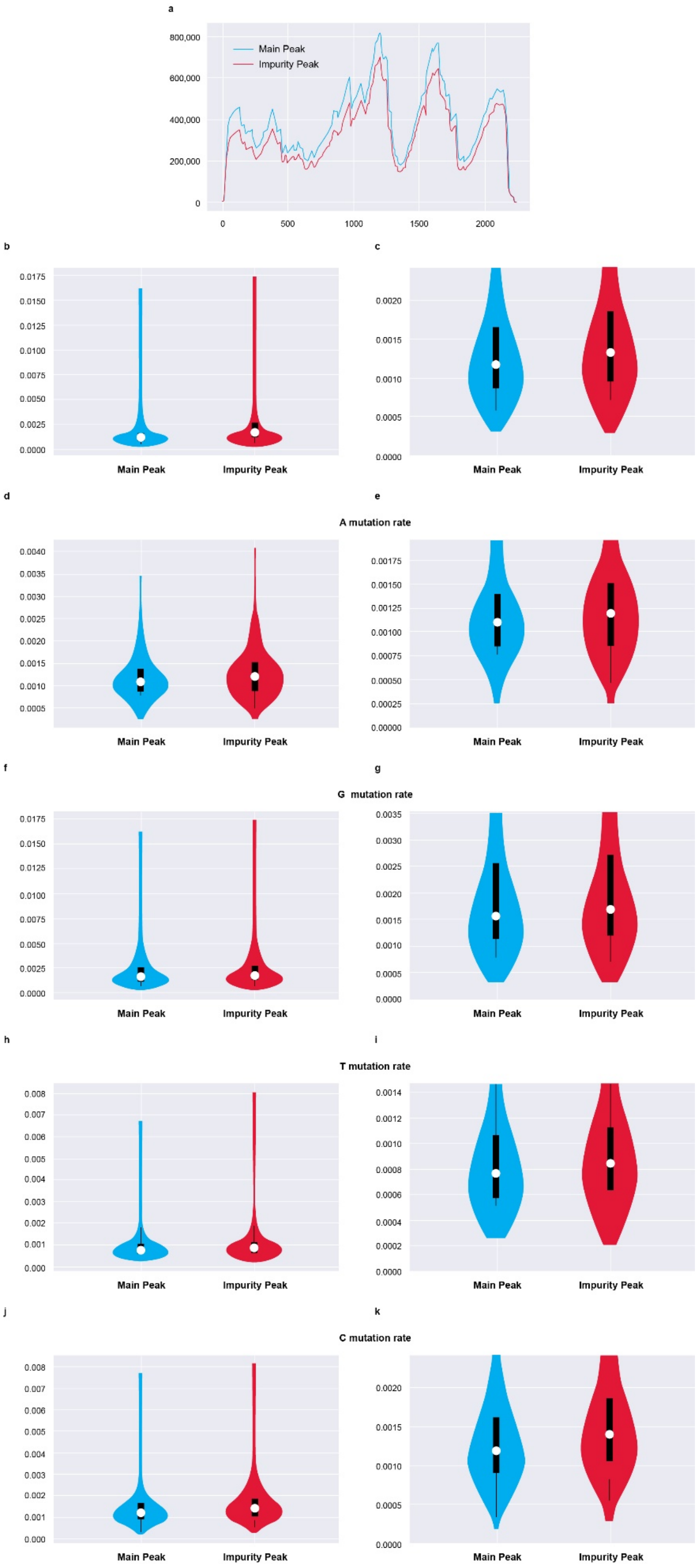
a
800,000
600,000
400,000
200,000
0
0
500
1000
1500
2000
Main Peak
Impurity Peak
b
c
d
e
A mutation rate
f
g
G mutation rate
h
i
T mutation rate
j
k
C mutation rate
Main Peak
Impurity Peak

**Supplementary Figure 3.** Isolated main peak (MP, blue) and late eluting-peak (LP, red) were sequenced by Illumina NGS. Box plots display median (white dot) and interquartile ranges (black box), and 1.5x the interquartile range in the whiskers. Distribution (maximum and minimum) is displayed as violin plots (blue and red), for two illumine squencing runs (read depth >10,000 reads per position). Read depth counts showed comparable sequence coverage across both fractions, and mutation rates both on the entire sequence and by individual bases showed no bias in the LP. These data failed to identify base-specific mutations or drops in sequence coverage that would indicate high levels or site-specific lipid modification of the nucleobases. Representative data from 2 repeat experiments are shown.

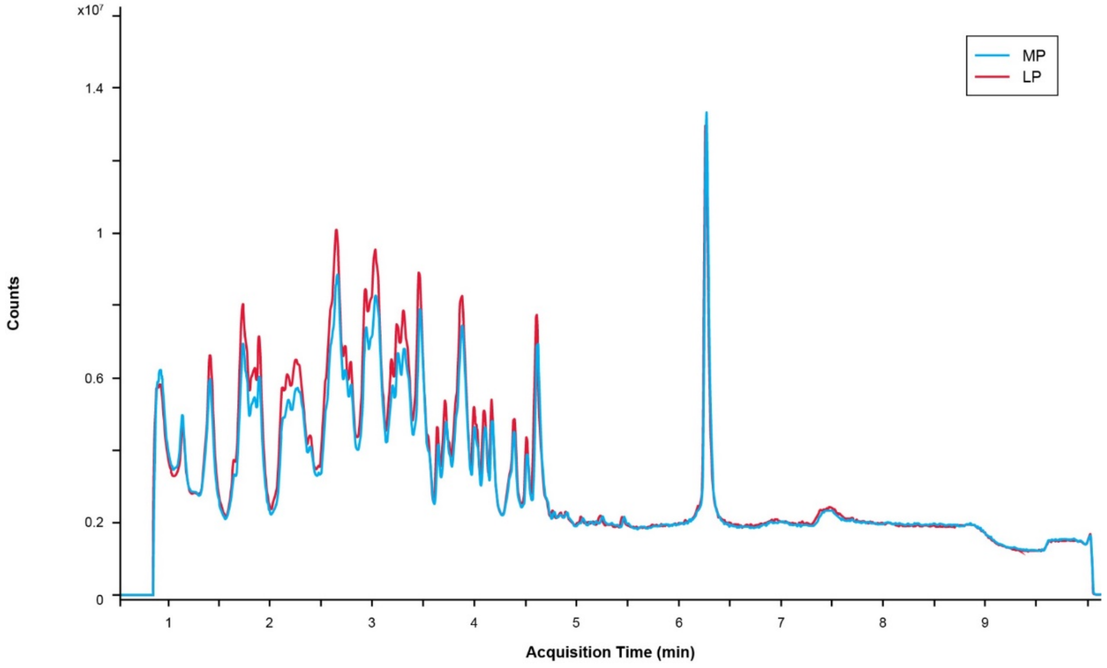


**Supplement Figure 4.** Isolated main peak (MP, blue) and late eluting-peak (LP,red) were subjected to T1 nuclease digest and analyzed by liquid chromarography/UV spectroscopy/mass spectrometry (LC/UV/MS) to yield an oligo fingerprint, as seen in the overlay. Overlaid total ion chromatograms (TICs) are shown in the chromatogram. Oligos corresponding to the entire sequence are resolved between 0.8 and 5.5 minutes, and the tail fragment at 6.3 minutes. The LP region showed the same sequence content as the MP. Additional study by differential analysis did not identify any differences. Representative data from 2 repeat experiments are shown.

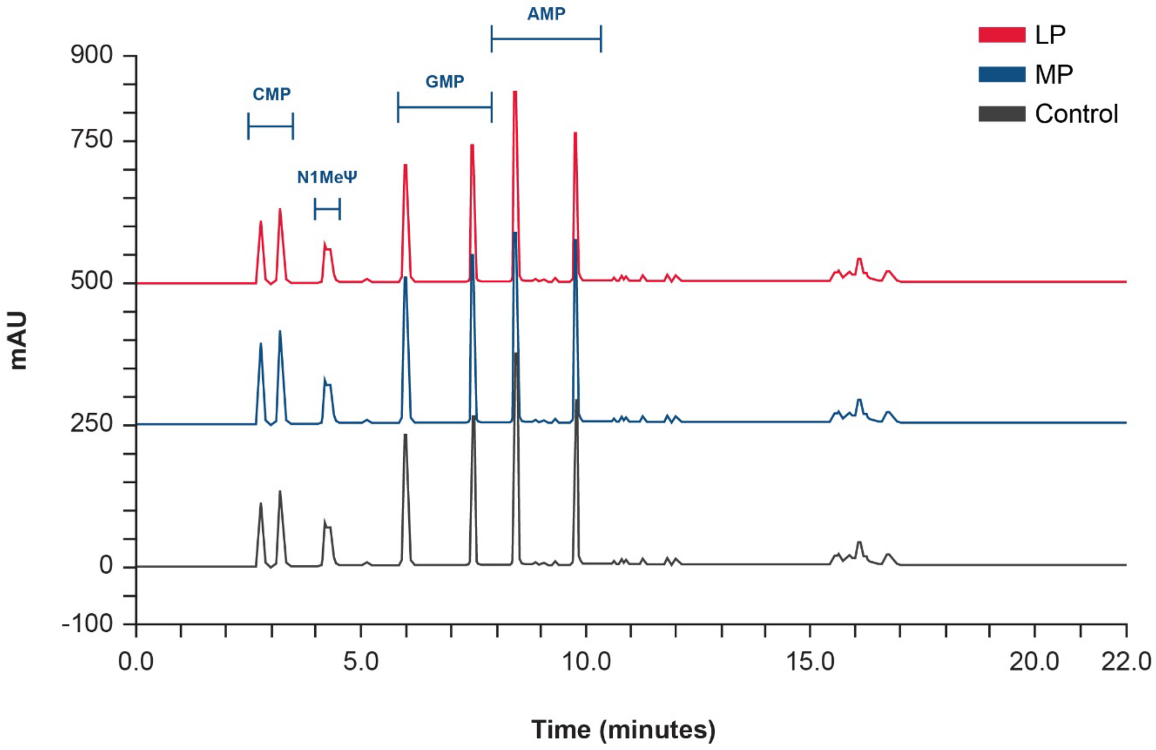


**Supplement Figure 5.** Isolated mRNA main peak (MP, blue), late eluting-peak (LP, red) and control (black) were subjected to total hydrolysis by 30-minute incubation at 65°C in 0.33 M NaOH, followed by neutralization with 1M HCl and ion-pair reversed phase liquid chromatography-UV (RP-IP LC-UV) analysis on a Waters BEH column with 100 mM TEAA and an acetonitrile gradient. Peaks were confirmed by 3D UV data. Complete digestion to nucleotide monophosphates was achieved, with racemic mixtures of 5` and 3` phosphate yielding two resolved peaks for each base. No difference in nucleotide content between the MP and LP was observed. Representative data from 2 repeat experiments are shown.

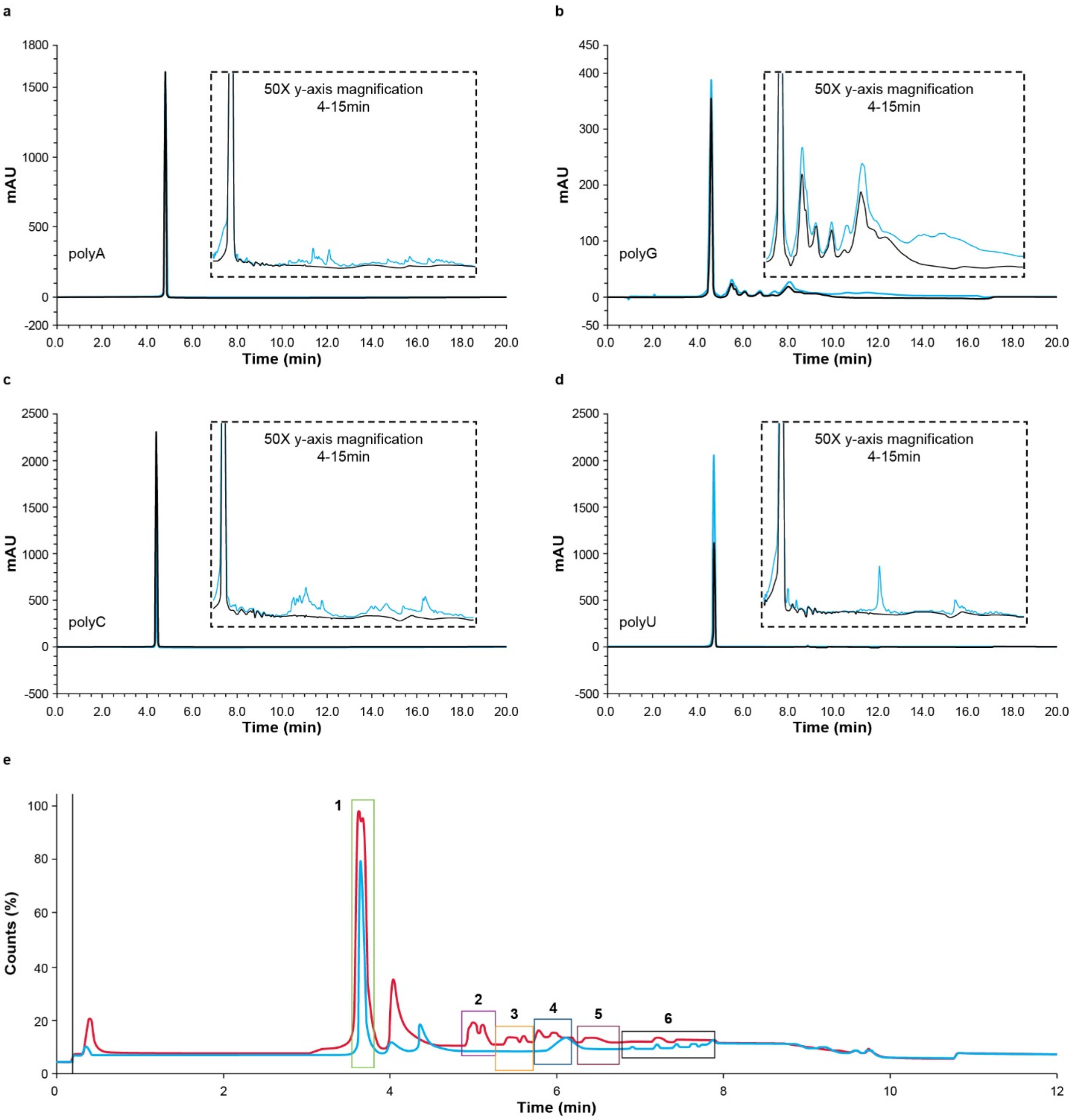


f

| Peak | Retention time (min) | Deconvoluted mass (amu) | Mass shift (amu) |
|---|---|---|---|
| 1 | 3.56-3.81 | 12226 | 0 |
| 2 | 4.91-5.20 | 12492 | 266 |
| 3 | 5.28-5.69 | 12604 | 378 |
| 4 | 5.70-6.24 | 12791 | 565 |
| 5 | 6.29-6.76 | 12903 | 677 |
| 6 | 6.77-7.84 | 13218 | 992 |

**Supplement Figure 6.** RNA homopolymers of adenosine (A), cytosine (C), guanosine (G), and urasil (U) of 40 nucleotides in length were individually prepared in the binary system with two ionizable lipid materials, incubated for 1 month at room temperature, and analyzed by liquid chromarography/UV/mass spectrometry (LC/UV/MS; n=1). **a-d**, Ion-pair reversed phase liquid chromatography-UV (RP-IP LC-UV) overlays of the pure oligo (black) and extracted binary (blue) are shown, with the unmodified oligo eluting between 4 and 5 minutes. Additional peaks in the polyG oligonucleotide in **(b)** are unsurprising and can be attributed to secondary/tertiary structure. Baseline magnification is inset for each trace, showing a polydisperse collection of lipid-modified peaks for all four nucleobases. **e**, The MS total ion chromatograms (MS TIC) is shown for polyC40 as an example with the pure oligo in blue and degraded ionizable lipid binary in red. Deconvolution was performed on the six boxed regions to obtain the data shown in (**f**). Peak 1 is the unmodified oligo, and subsequent retention times show an increase in mass attributed to addition of a lipid chain. Comparable masses were observed for other nucleobases, consistent with addition of a fragment of the ionizable lipid. AU, absorption units.

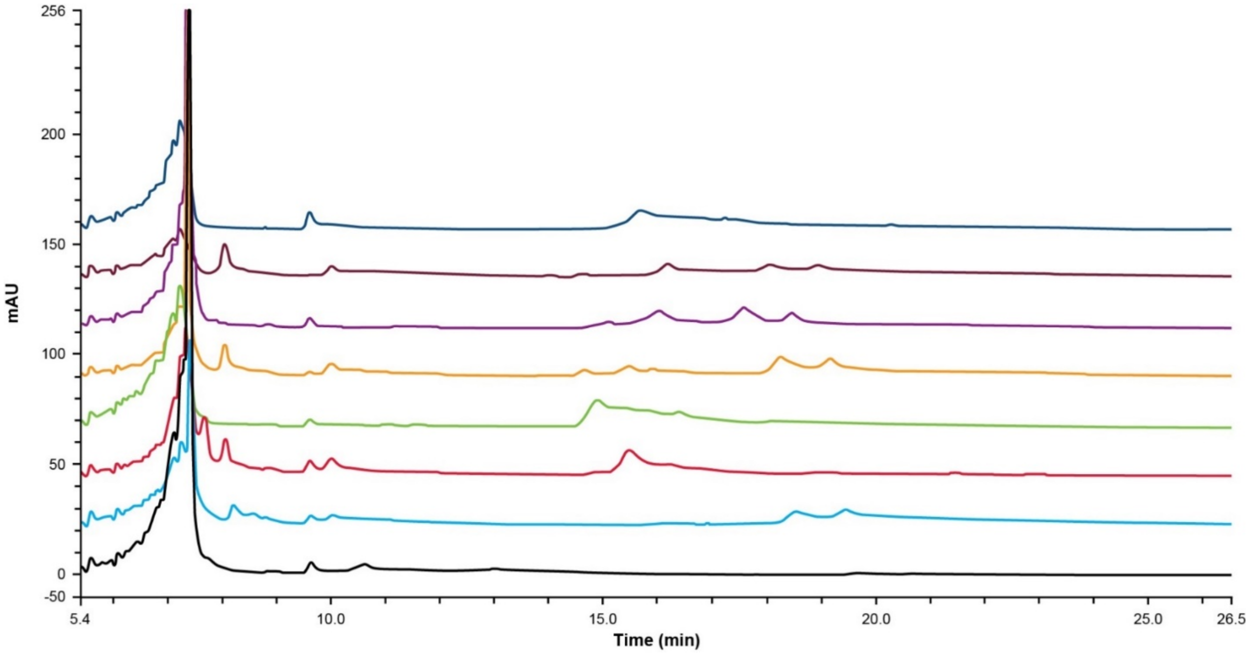


**Supplement Figure 7.** Eight different ionizable lipid chemistries were prepared in binaries with mRNA. Reversed phase-ion pair (RP-IP) chromatograms of the extracted mRNA are shown, with the unmodified mRNA eluting at 7 minutes, and lipid-modified mRNA eluting from 9 to 25 minutes. Each color represents a different ionizable lipid tested (identities of lipids are proprietary). Each ionizable lipid resulted in a different peak profile, with retention times of the lipid modified species corresponding to different resulting hydrophobicities based on lipid modification. 7 representative chromatograms from >100 lipid chemistries analysed are shown.

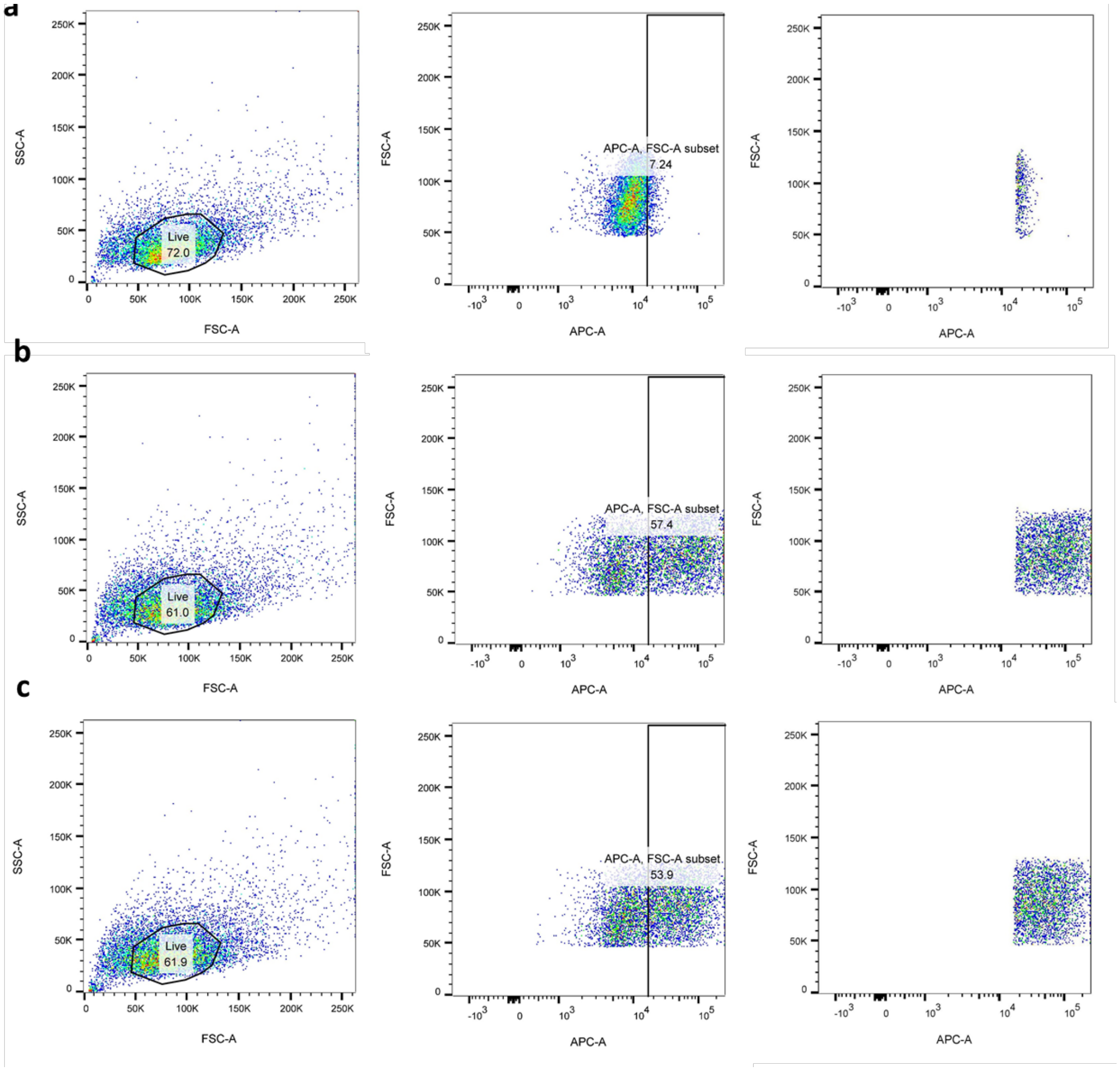


**Supplement Figure 8.** Gating strategy for detecting protein expression in HeLa cells by flow cytometry for **a** negative mRNA control, **b** positive mRNA control, and **c** representative lipid nanoparticle (LNP) sample. Cells with detectable Allophycocyanin-A (APC-A) conjugated proprietary antibody fluorescence ≥5% of the non-translating mRNA negative control were considered to be positive.
